# Perinatal environmental enrichment affects murine neonates’ brain structure before their active engagement with environment

**DOI:** 10.1101/2025.02.22.639627

**Authors:** Malte S. Kaller, Clémence Ligneul, Rylan Allemang-Grand, Tie Yuan Zhang, Jacob Ellegood, Michael Meaney, Jason P. Lerch

**Affiliations:** Wellcome Centre for Integrative Neuroimaging, FMRIB, Nuffield Department of Clinical Neurosciences, University of Oxford; Mouse Imaging Centre, The Hospital for Sick Children, Toronto, Ontario, Canada; Department of Medical Biophysics, University of Toronto, Toronto, Ontario, Canada; Douglas Hospital Research Centre, Department of Psychiatry, McGill University, Montréal, Canada; Bloorview Research Institute, Holland Bloorview Kids Rehabilitation Hospital, Toronto, Ontario, Canada; Institute for Human Development and Potential (IHDP), Agency for Science, Technology and Research (A*STAR), Singapore, Singapore; Douglas Mental Health University Institute, McGill University, Montreal, QC, Canada; Program in Child and Brain Development, CIFAR, Toronto, ON, Canada

## Abstract

Early life experiences shape individuals. Environmental enrichment, an experimental paradigm used to study the effect of increased environmental complexity and novelty in animal models, has long been recognised for its broad effect on nervous system function and behaviour. In adult rodents, structural changes in the brain due to enriched environments are well documented, notably in the hippocampus. However, the effect of environmental enrichment on the developing brain during early life is not well understood. This study aims to investigate how environmental enrichment affects brain development during the critical perinatal period, and how such effects compare to those observed during adulthood.

We use high-resolution MRI to measure the brain structure of mouse neonates at postnatal day 7, born either in an enriched or a standard environment. We show that rodents exhibit brain structure differences as early as postnatal day 7. However, the regional changes observed differ from those in adulthood: hippocampal changes are limited, but changes in the hindbrain, the dorsal striatum, and the medial habenula are strong.

Given the lack of direct interaction between neonates and the environment at P7, we hypothesised that maternal care may mediate these effects. We show that maternal care differs between enriched and standard environments, that maternal care correlates with brain structure changes in the neonates, and that maternal care and enriched environment affect brain structure similarly. Further analysis suggests that early changes in brain structure due to environmental enrichment are at least partly mediated by maternal care. This study provides novel insight into the differential effect of enriched environment on early brain development in rodents.

## Introduction

Environmental enrichment (EE) is an experimental paradigm used to study the effects of increased environmental stimulation in animals. It can encompass aspects such as enhanced spatial navigation, social contact, exercise, novelty, and overall more complex housing conditions. This increased stimulation can induce various plastic responses in the adult brain, including changes in biochemical parameters, dendritic arborization, neurogenesis, and enhanced learning (van Praag et al., 2000). In adult mice, exposure to a novel, enriched environment leads to rapid structural brain changes detectable via MRI, with the hippocampus being particularly sensitive to EE (Scholz et al., 2015).

While the effects of EE on adult brain plasticity are well-characterized, much less is known about its influence during early development. However, the brain is highly sensitive to environmental stimulation during early critical developmental windows, and changes in brain development during this period may have long-term consequences (Reh et al., 2020). Efforts to study EE in early life focus mostly on the visual system (Baroncelli et al., 2010), reproducibly showing an accelerated maturation of the latter and an increase in BDNF and IGF-I expression in neonate brains. Other work focused on the effect of early development when the dam is exposed to enrichment during gestation. Animals born from a dam exposed to a free running wheel during gestation display lower weights from P8 to P49, but higher cell proliferation in the dentate gyrus at P8 (Bick-Sander et al., 2006). Additionally, neonates from dams exposed to EE before birth (Cárdenas et al., 2015) have an improved basic behaviour (e.g., improved righting reflex, gait and negative geotaxis). These studies indicate that early changes in environment might affect brain organisation, even before the neonates can explore their environment.

How EE drives plastic changes in the brain during early development is still an open question. Dams exposed to an enriched environment can exhibit different maternal behaviours (Sparling et al., 2020), which could act as a catalyst for the effects of EE on neonates. Additionally, pioneering studies show that maternal behaviour affects the neonates glucocorticoid system and drives epigenetic changes in brain regions, including the hippocampus (Meaney et al., 2007; Turecki and Meaney, 2016). However, the effects of perinatal EE on whole-brain structure and organisation remain poorly understood, and the potential mediating role of maternal care during early development remains hypothetical.

This study aims to understand the effects of environmental enrichment (**Figure 1A**) on brain structure as an intrinsic environmental factor present from birth (**Figure 1B**, dataset P). Additionally, we investigate how the effects of perinatal enrichment compare to those of enrichment provided only in adulthood (**Figure 1B**, dataset A). First, we replicate previous findings showing a strong hippocampal response to enrichment in adulthood. We then identify brain structures responding differently depending on whether animals were born into EE or introduced to it later in life. To explore the origins of these differences, we examine the effects of enrichment at postnatal day 7 (P7), before neonates can actively explore their surroundings (**Figure 2**, dataset N). We hypothesise that the dam’s exposure to EE influences maternal care, thereby affecting neonatal brain structure. Our findings reveal significant structural differences in the brains of neonates (P7) exposed to perinatal EE and provide data supporting the role of maternal care in mediating these effects.

**Figure 1:**
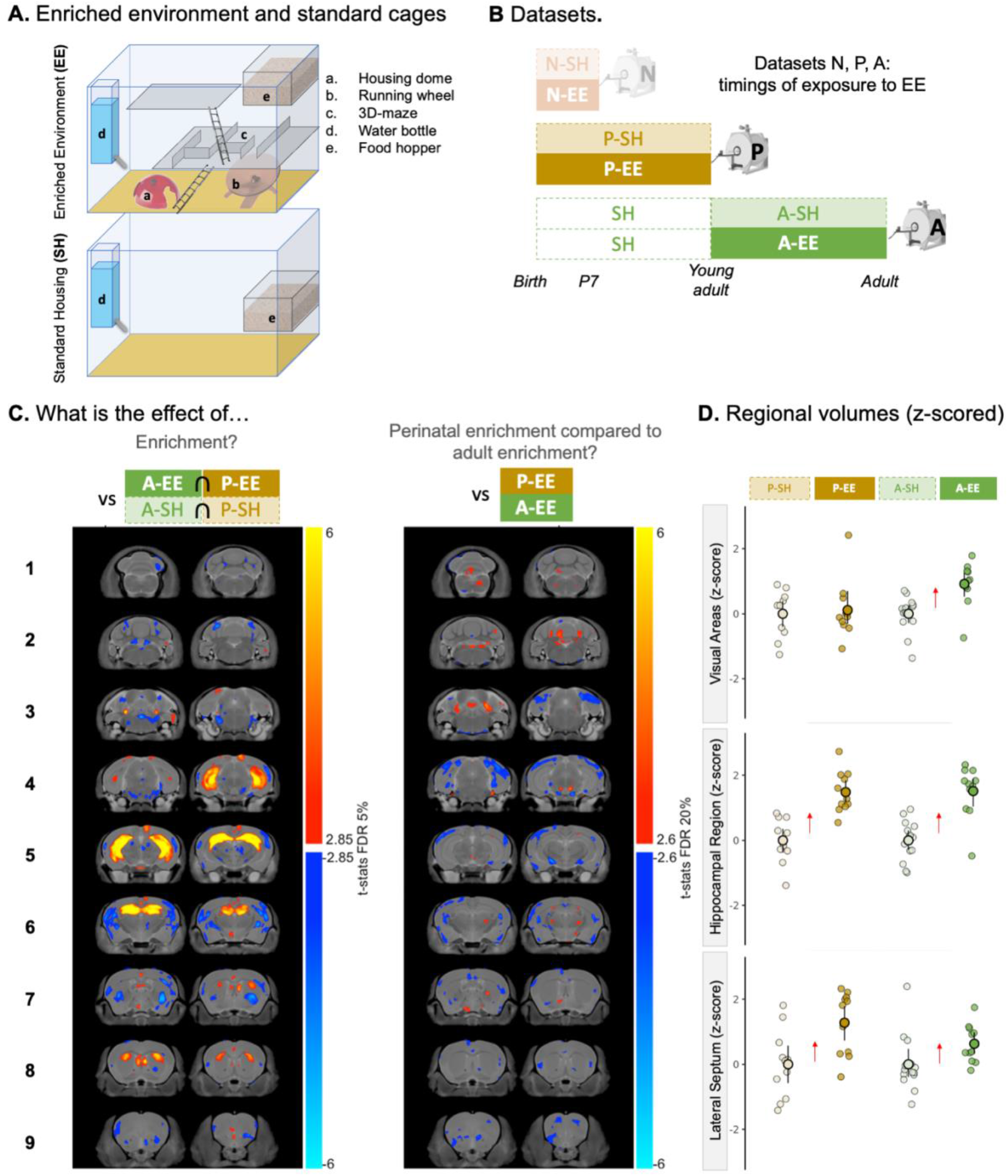
Effect of perinatal and adult exposure to enrichment observed in adulthood. **(A)** Schematics of the enriched environment (EE) and Standard Housing (SH). **(B)** Illustration of the datasets used in the paper. Dataset N (“neonatal”): Perinatal enrichment, neonates perfused at P7 for ex vivo MRI. N-EE: Neonates born in EE (dams in EE from E13); N-SH: Neonates born in SH. Shaded because not used in this figure. Dataset P (“perinatal”): Perinatal enrichment until adulthood (6 weeks of enrichment), animals perfused at P43 for ex vivo MRI. P-EE: Animals born in EE (dams in EE from E17). P-SH: Animals born in SH. Dataset A (“adulthood”): Animals in standard housing until P53, adulthood enrichment from P53 to P96 (6 weeks of enrichment). Animals perfused at P96 for ex vivo MRI. A-EE: Animals transferred to EE in adulthood. A-SH: Animals staying in SH in adulthood. More details are provided in the Methods section. **(C)** A voxel-wise linear model was applied to the Jacobians computed after linear co-registration (hence corrected for individual brain volume variation) coming from datasets P and A (see Methods). (left panel) Effect of EE in adulthood regardless of the timing of enrichment. The regressors are housing condition, age and sex. (right panel) Differential effect of perinatal vs adulthood enrichment. The regressors are housing, age and sex, and the interaction term housing*age. **(D)** z-scored regional volumes, normalised by total brain volume (only male volumes shown to address the sex discrepancy between datasets P and A). Data are zeroed to the mean of the standard housing condition in their age group. The z-score is calculated using the mean and the standard deviation is shared over datasets P and A. The error bars represent the lower and upper confidence limits.

**Figure 2:**
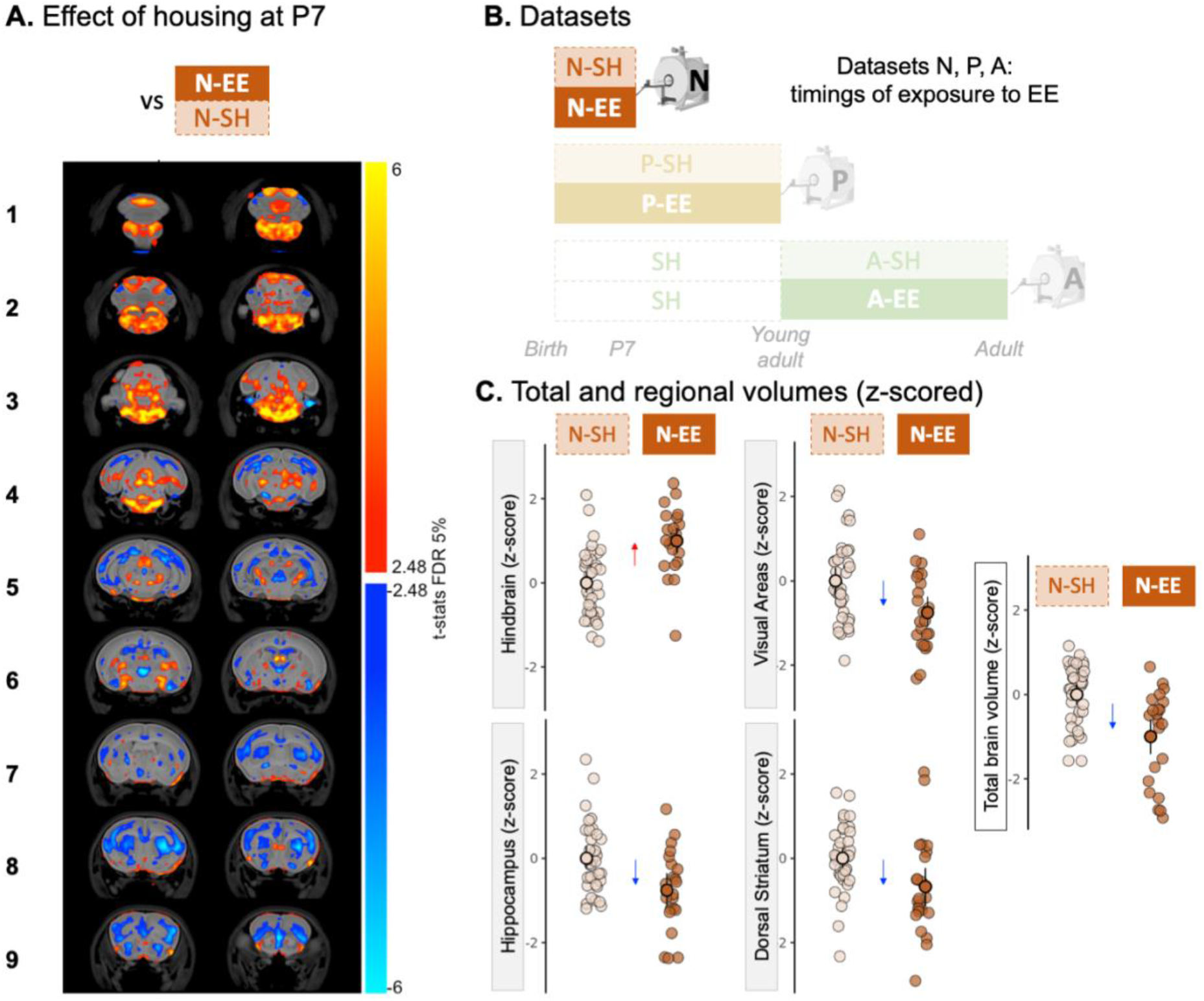
Effect of perinatal exposure to enrichment observed at P7. **(A)** Effect of EE in neonates (P7). A linear model was applied to the Jacobians computed after linear co-registration (hence corrected for individual overall brain volume variation), with the following housing condition, litter order, litter size and sex as regressors. The effect of housing condition is shown with a FDR threshold at 5%. **(B)** Datasets of the paper with timings of exposure (c.f. caption Figure 1B). Data in this figure come from the dataset N. **(C)** z-scored regional volumes normalised by total brain volume, and z-scored total brain volumes (only male volumes shown to be consistent with datasets shown in Figure 1D). Data are zeroed to the mean of the standard housing condition. The z-score is calculated using the mean. The error bars represent the lower and upper confidence limits.

## Results

We analysed three datasets with different timings of exposure to an enriched environment (C.f. Methods – Enrichment Setup). The adult housing dataset (A) assessed the effects of enrichment introduced in adulthood, where mice were exposed to enriched conditions from P53 to P96. The perinatal-to-adult housing dataset (P) included mice reared in enriched or standard environments from late gestation (E17) until P43. Exposure duration to enrichment was the same between datasets A and P. The P7 perinatal housing dataset (N) examined neonates at postnatal day 7 (P7) whose dams were housed in either enriched or standard conditions from embryonic day 13 (E13) (c.f. Methods – Animals and Methods – Datasets for more details).

To understand the impact of environmental enrichment on the brain in each condition (dataset A, P and N), we used ex vivo high-resolution magnetic resonance imaging (MRI) (c.f. Methods – MRI Scanning). Brain samples were then coregistered (dataset A and P together, dataset N separately), generating deformation maps, and therefore providing high sensitivity to structural differences between animals and groups (c.f. Methods – Analysis / Image Registration). Results are presented either whole brain t-stats maps (coming from voxel-based morphometry) showing overall volume differences, or regional volume differences. Maternal care was assessed before the acquisition of dataset N (c.f. Methods - Behaviour).

### Neuroanatomical signature in adulthood of long-term exposure to environmental enrichment on brain volumetric changes: the hippocampus is the primary focus, regardless of the timing of enrichment

Animals exposed to EE during 43 days, either from birth or in adulthood (n=37, CD1 mice, A-EE and P-EE), were compared to animals in standard housing (SH) (n=39, CD1 mice, A-SH and P-SH) using volumetric analysis of T2-weighted ex vivo MRI data (see Methods – MRI Scanning). Animals exposed to EE exhibit a strong hippocampal volume increase (voxel-based analysis, **Figure 1C, left panel**; regional analysis Cohen’s d=1.48 for dataset P and Cohen’s d =1.51 for dataset A, **Figure 1D, middle panel**). Bilateral volume increases are also observed in the lateral septum complex (Cohen’s d =1.28 for dataset P and Cohen’s d =0.63 for dataset A, **Figure 1D, lower panel**), the dorsal-rostral caudoputamen, and cerebellar lobules IV-V. Bilateral volume decreases are observed in the ventral-caudal caudoputamen, secondary somatosensory areas, fourth ventricle and cerebellar lobule VI. White matter tracts also show volumetric increase (fimbria) and decrease (middle cerebellar peduncle). These observations result from a linear model accounting for housing condition, age and sex that was applied to each voxel, and the resulting t-stats map was thresholded at 5% FDR. Considering the male/female ratio discrepancy between the perinatal and adult enrichment datasets (see **Supplementary Table 1** for details), we ran the same analysis with the males only. Results are similar, but less powered (See **Supplementary Figure 1**). The regional volume differences (**Figure 1D**) are shown in males only to facilitate the effect-size estimation, as brain maturation is sexually dimorphic (Qiu et al., 2018; Juraska and Drzewiecki, 2020).

Overall, EE reproducibly affects brain structure in adulthood, notably the hippocampus and a few other subcortical regions, confirming previous results (Scholz et al., 2015). The strength of the effect on hippocampal volumes is very similar if enrichment is started during adulthood, whilst for lateral septum complex volumes, the effect is stronger if enrichment is present at birth. As discussed in the next section, the regional effect size sometimes depends on when enrichment was introduced.

### Neuroanatomical signature of perinatal enrichment: frontal regions, sensory areas and ventral tegmental area (VTA) are differentially affected by the timing of exposure to environmental enrichment

Although most structural effects of enrichment are similar when animals are exposed perinatally or during adulthood, a few regions are differentially affected by the timing of the enrichment. A voxel-wise linear model was applied, accounting for the interaction term between housing condition and age plus sex (**Figure 1C, right panel**). An atlas segmentation was used to retrieve regional volumes (see Methods). Several cortical regions are typically smaller under EE in animals that were exposed to EE perinatally compared to animals that were exposed in adulthood.

These regions comprise notably the olfactory areas, and more specifically the piriform area (Cohen’s d=-1.26 for dataset P and Cohen’s d=0.70 for dataset A), the auditory areas (Cohen’s d=-0.95 for dataset P and Cohen’s d=0.57 for dataset A), the somatomotor areas (Cohen’s d=-0.75 for dataset P and Cohen’s d=0.21 for dataset A), the orbital area (Cohen’s d=-0.81 for dataset P and Cohen’s d=0.59 for dataset A). The visual areas are stable in animals that were exposed to EE perinatally, but they are bigger in animals that were exposed to EE in adulthood (Cohen’s d=0.11 for dataset P and Cohen’s d=0.92 for dataset A, **Figure 1D, upper panel**). On the other hand, some regions are relatively larger in animals that were exposed to enrichment perinatally, such as the cerebellar lobules IV-V (Cohen’s d=1.05 for dataset P and Cohen’s d=-0.55 for dataset A). Interestingly, as can be seen on voxel-based maps, a few thalamic nuclei, and most notably the ventral tegmental area (VTA), are also larger in animals that received perinatal enrichment.

These differences suggest different mechanisms in the brain’s response to enrichment if animals are born in an enriched environment or if their environment is enriched in adulthood.

### Brain structure is strongly impacted by the enriched environment at P7, with a different pattern compared to adulthood

To understand the origins of the differences between the adult and perinatal enrichment effects (i.e., datasets P vs A, **Figure 1C, right panel**), we measured brain structure in neonates at P7 born either in enriched or standard environments (dataset N), when they were not able to explore their surroundings yet. Dataset N consists of 5 dams, whose neonates were either born in EE or SH environments. Dams were mated a second time, and 2 of them switched environments compared to the first experiment, resulting in a total of 119 neonates (52 animals born in EE, see **Methods** for more details). We observe volumetric brain differences between both environments at P7 (**Figure 2**). Because the T1/T2 contrast in the neonates is very poor, the images were acquired with a diffusion-weighted sequence to improve contrast in the neonatal brain.

First, we report an overall brain size difference between the neonates born in the enriched environment (264±23µl, Cohen’s d =-1.00) compared to those born in the standard environment (289±18µl) (**Figure 2C**). Note the very high number of neonatal brains scanned (118) compared to datasets A and P, yielding highly significant results.

Then, a linear model was applied voxel-wise to the Jacobian maps computed after linear co-registration (hence corrected for individual brain volume variation), with housing condition (EE vs SH), litter order (dam’s first litter vs dam’s second litter), litter size and sex as regressors. We report a consistent effect of housing condition in the frontal regions of the brain (volume decrease in frontal lobe & striatum), in a few thalamic nuclei (notably volume increase in medial habenula), and in the brain stem (volume increase pons and medulla). The atlas segmentation at P7 offers a coarser precision than the atlas used in adulthood, but we could retrieve effect sizes for some ROIs. The volume decrease is important in the striatum (Cohen’s d=-0.67), in the parietal lobe, containing some auditory, somatosensory and somatomotor areas (Cohen’s d =-0.90) and in the occipital lobe, assimilable to visual areas (Cohen’s d =-0.76). The hippocampus (Cohen’s d =-0.76) is also affected, but note the opposite direction of change compared to observations in adulthood. A strong volume increase is noted for the hindbrain (Cohen’s d =1.00). The effect of litter order and the effect of litter size are present, but they are considerably sparser than the effect of enrichment (**Supplementary Figure 3**).

The pattern of volumetric changes due to perinatal enrichment is different at P7 and adulthood, as can be seen in **Supplementary Figure 4** (N, P, A and A|P comparison) and more precisely reported in **Supplementary Table 2**. Although the brain organisation is overall similar at birth and in adulthood, one-to-one comparisons are not always possible (notably because of segmentation). The identifiable regions that present similar changes due to perinatal enrichment at P7 and in adulthood are the visual areas, potentially some thalamic nuclei and some cerebellar lobules. The volume of ventral caudoputamen is reduced regardless of the moment when the effect of enrichment is observed or introduced, and the hippocampus is affected at all ages, although the volumetric change at P7 is opposite to the volumetric change observed in adulthood.

Particular attention should be brought to the difference in litter sizes between dams’ first litter and dams’ second litter (**Supplementary Table 1**). The litter size of the second litter was controlled, being kept at 11 neonates per dam. The first litters were not controlled for litter size: note the extremely large litter size of Cage A (17 neonates). Litter size notably affects neonates’ birth weight and milk intake, and is a potential modulator of the amount of maternal care each neonate can receive.

### The effect of environmental enrichment on maternal behaviour

EE influences the brain volume of P7 neonates (**Figure 2**). However, mice at P7 show very little active engagement with their environment. Hence, it is not clear through which route EE may influence brain development: the dam could be a link. If the environment changes the dam’s behaviour and interaction with the neonates, their experience in very early life could be significantly different. Hence, we tested the hypothesis that EE will change maternal behaviour.

Assessment of maternal behaviour took place on P3 for 35 mins (between 10am and 12pm) and on P4 for four 30 mins sessions spread out between 9am and 5pm (total of 2 hours and 35 mins per cage) (see **Figure 3A**, or **Supplementary Figure 5A** for more detailed schematic design). This protocol for assessing maternal behaviour was based on Myers et al. (1989) and Champagne et al. (2003) with an emphasis on active nursing postures and grooming bouts. The total time dams spent in contact with the neonates (maternal contact time) was observed, as well as the time spent performing various behaviours while in contact with the neonates (see **Supplementary Table 3** & method section for more details).

**Figure 3:**
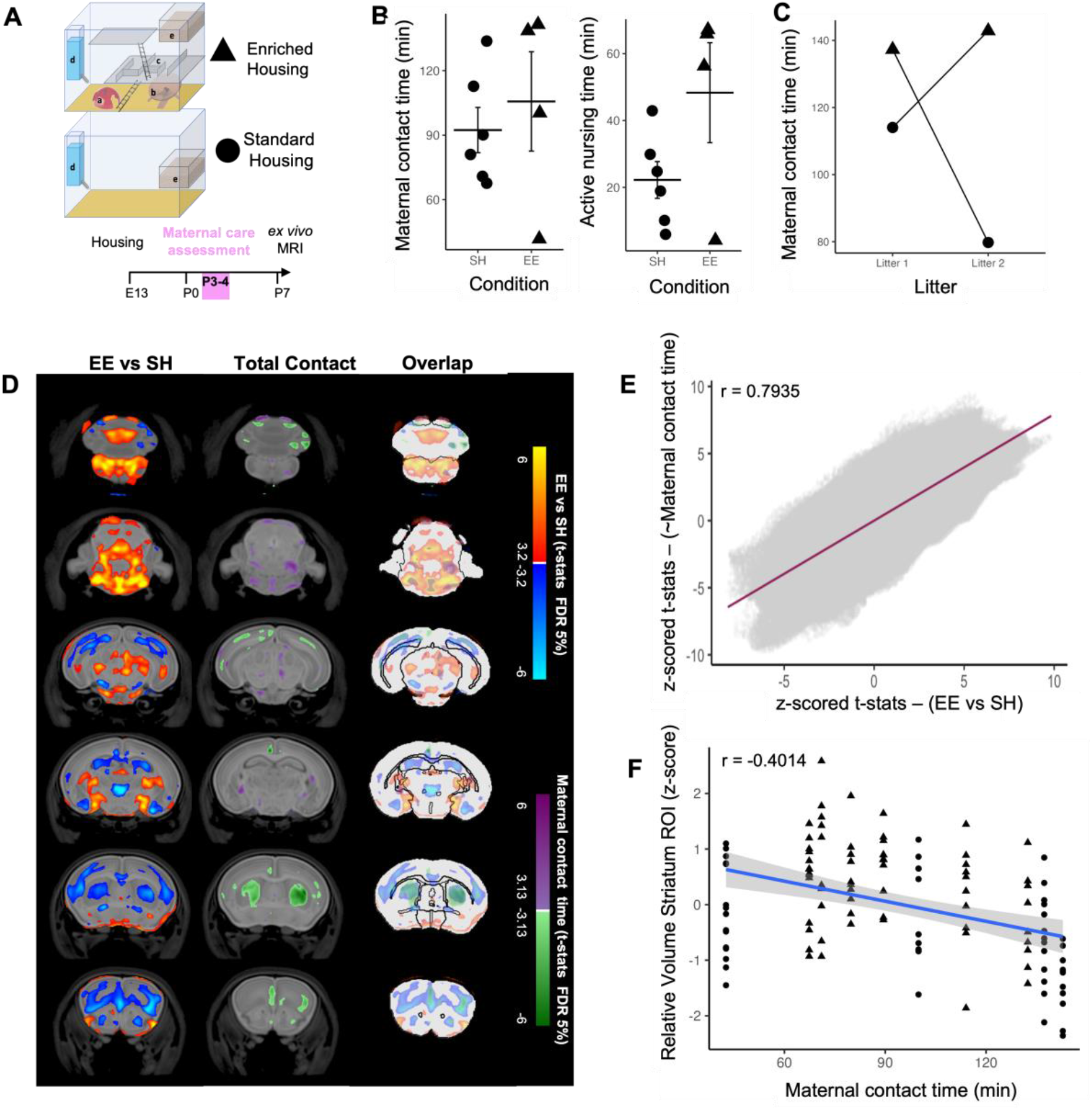
Effect of housing on maternal behaviour and its relation to P7 brain structure. **(A)** Schematic experimental design. **(B)** Maternal contact time and time spent actively nursing the offspring per condition for each observation. C. Maternal contact time for dams that changed housing conditions between Litter 1 and Litter 2. **(D)** Voxel-wise t-statistics of the effect of housing (EE vs. SH, left) and maternal behaviour (Total Contact, middle) on brain volume, as well as both contrast overlayed on top of each other to emphasize similarities. **(E)** Z-score of the t-stats for each voxel from the voxel-wise brain analysis for housing and maternal behaviour plotted against each other and showing a strong correlation. **(F)** In an exemplary striatal ROI, in which enrichment led to a decreased volume, higher contact time of dams with their neonates predicted lower volume size (r= -0.4014).

*Maternal behaviour revealed overall higher metrics in EE compared to SH (Figure 3B). Notably, in dams in which housing condition was changed from the first to the second litter, the environmental change coincided with increased maternal contact time in EE vs SH (Figure 3C). However, the variability and the low sample size (EE, n=4 dams; SH, n=6 dams; Supplementary Figure 5 C&D) prevent a definitive conclusion on the influence of the environment on maternal care*. Maternal contact time is related to volumetric changes in P7 neonates

As EE seems to affect maternal behaviour, we tested whether the measured maternal behaviour influenced brain structure at P7. We hypothesised that maternal contact time, a coarse but robust measure of maternal behaviour, would significantly affect brain structure at P7.

We tested for the effect of maternal contact time on volumetric changes using a voxel-wise analysis, controlling for sex and the effect of litter size (**Figure 3D, middle**). The clusters (and their directionality) overlap clearly between the effect of environmental enrichment (**Figure 3D, left column**) and the effect of maternal behaviour (**Figure 3D, middle column**).

This overlap is further emphasised when plotting against each other the z-score of the t-stats for each voxel from the voxel-wise brain analysis (**see Figure 3E**). Overall, this indicates a significant correlation (r=0.79) between the effect of environmental enrichment and maternal behaviour on neonates’ brain development. Furthermore, time spent in contact with the neonates significantly predicts the volume of key brain areas, such as the striatum (**Figure 3F**) or brain stem (**Supplementary Figure 5F**).

### Mediation analyses of housing and maternal behaviour effects on brain structure

To test whether maternal care mediated the effects of housing condition (SH vs. EE) on brain structure at P7 (dataset N), we performed mediation analyses at both the region-of-interest (ROI) and voxel-wise levels (see Methods).

ROI-based mediation analyses demonstrated a strong effect of housing condition on striatal volume (total effect estimate = −1.04, 95% CI: −1.38 to −0.70, p < 0.001). Mediation analysis indicated that this effect was partially mediated by maternal contact behaviour: the indirect (mediated) effect was significant (ACME = −0.47, 95% CI: −0.77 to −0.19, p = 0.0004), while the direct effect of housing condition remained significant after accounting for maternal contact (ADE = −0.57, 95% CI: −0.98 to −0.17, p = 0.0044). Approximately 45% of the total effect of housing on striatal volume was mediated by maternal contact (95% CI: 18% to 79%). In contrast, mediation effects in the brainstem ROI were weaker and less consistent. ROI-level mediation effects were assessed using both frequentist and Bayesian multilevel approaches, which yielded qualitatively similar patterns, supporting the robustness of the observed striatal mediation despite the distinct assumptions of each method (see Methods).

To explore further the spatial distribution of maternal care–mediated housing effects across the brain, we performed a voxel-wise mediation analysis to examine the directionality and relative strength of total, direct, and indirect effects of housing on brain volume (**Figure 4**). Due to the substantial computational demands of voxel-wise modelling, this analysis was performed using a frequentist mediation framework. The total effect maps revealed widespread housing-related structural differences (**Figure 4A**). Proportional mediation map (ACME / total effect) suggested regionally specific mediation by maternal care (**Figure 4B**). Decomposition of these effects in ROI indicated prominent direct effects (ADE) of environmental enrichment on brainstem volume (**Figure 4C**), with while indirect mediation effects of maternal care are notable effects in striatal regions (**Figure 4D**). Across the brain, the relative contribution of direct and indirect pathways varied spatially, highlighting regional heterogeneity in the mechanisms linking housing environment to brain structure. These voxel-wise results are descriptive and exploratory in nature.

**Figure 4.**
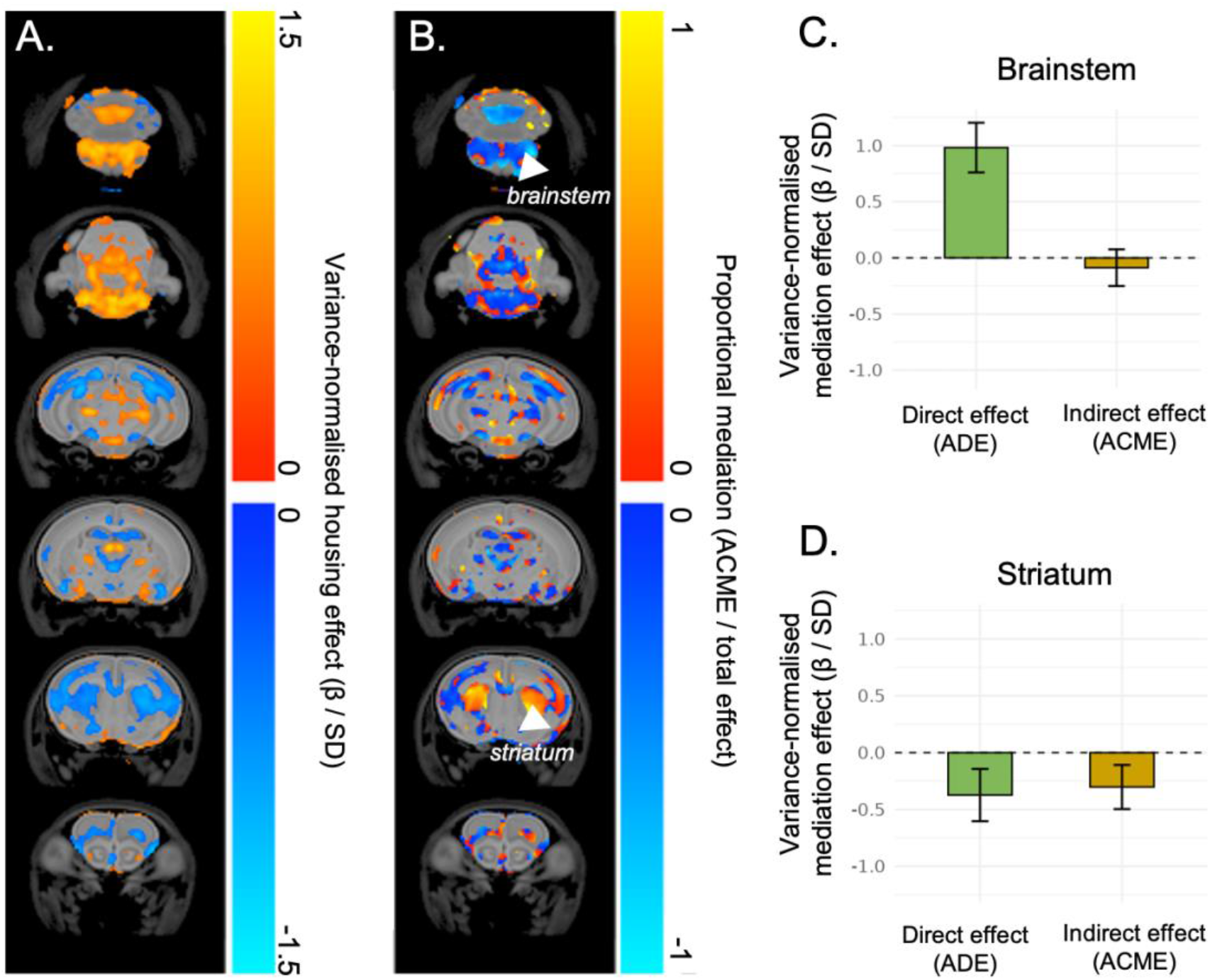
Voxel-wise mediation effects of housing on brain structure. **(A)** Voxel-wise total effect of housing on relative Jacobian values, expressed as a variance-normalised effect (β / SD), where β is the model-estimated effect size and SD is the voxel-wise inter-subject standard deviation. This scaling provides a dimensionless measure of effect magnitude relative to natural anatomical variability. **(B)** Proportional mediation map (ACME / total effect), showing the fraction of the housing effect statistically attributable to maternal care. Warm colours indicate positive proportional mediation (indirect and total effects in the same direction), whereas cool colours indicate negative proportional mediation (indirect and total effects in opposite directions). **(C–D)** Region-of-interest (ROI) summaries of direct and indirect mediation effects in the brainstem **(C)** and striatum **(D).** Bars represent variance-normalised effect estimates (β / SD) for the direct effect (ADE: Average Direct Effect of housing on voxel values) and indirect effect (ACME: Average Causal Mediation Effect via maternal care). Error bars denote ± SD across voxels within each ROI. ROI masked extracted from housing effect (see method). Mediation analyses were performed voxel-wise using linear models (mediator model: Total Contact ∼ Housing + Size Litter; outcome model: voxel value ∼ Housing + Total Contact + Size Litter).

## Discussion

We explored how brain structure responds to being born in an enriched environment (EE). First, we replicated the effects of environmental enrichment (EE) on brain structure in adulthood (Scholz et al., 2015). Furthermore, we found sparse structural differences between animals born in an enriched environment (EE) and those transferred to an enriched environment (EE) in adulthood after 6 weeks of exposure (A-EE vs P-EE, **Figure 1C**, right panel). Lastly, we provided novel evidence that animals born in EE exhibit strong differences in brain structures as early as P7 compared to animals raised in standard housing (N-EE vs N-SH, **Figure 2A**). Unexpectedly, the affected regions differ largely from those identified in adulthood or in the differential maps (A-EE vs. P-EE). This suggests the prominent condition-induced changes at P7 are partly transient, with limited influence on brain volume in later life. The experimental design included monitoring of maternal behaviour. While evidence of an environmental effect on maternal behaviour is only suggestive, we found that maternal behaviour was associated with structural changes in P7 neonates in regions overlapping with the environmental effects. Overall, our data provide evidence that an enriched environment influences brain structure in mice as early as P7, which may at least partially be mediated by differences in maternal care as suggested by the exploratory mediation analysis **(Figure 4**, **Figure 5).**

**Figure 5:**
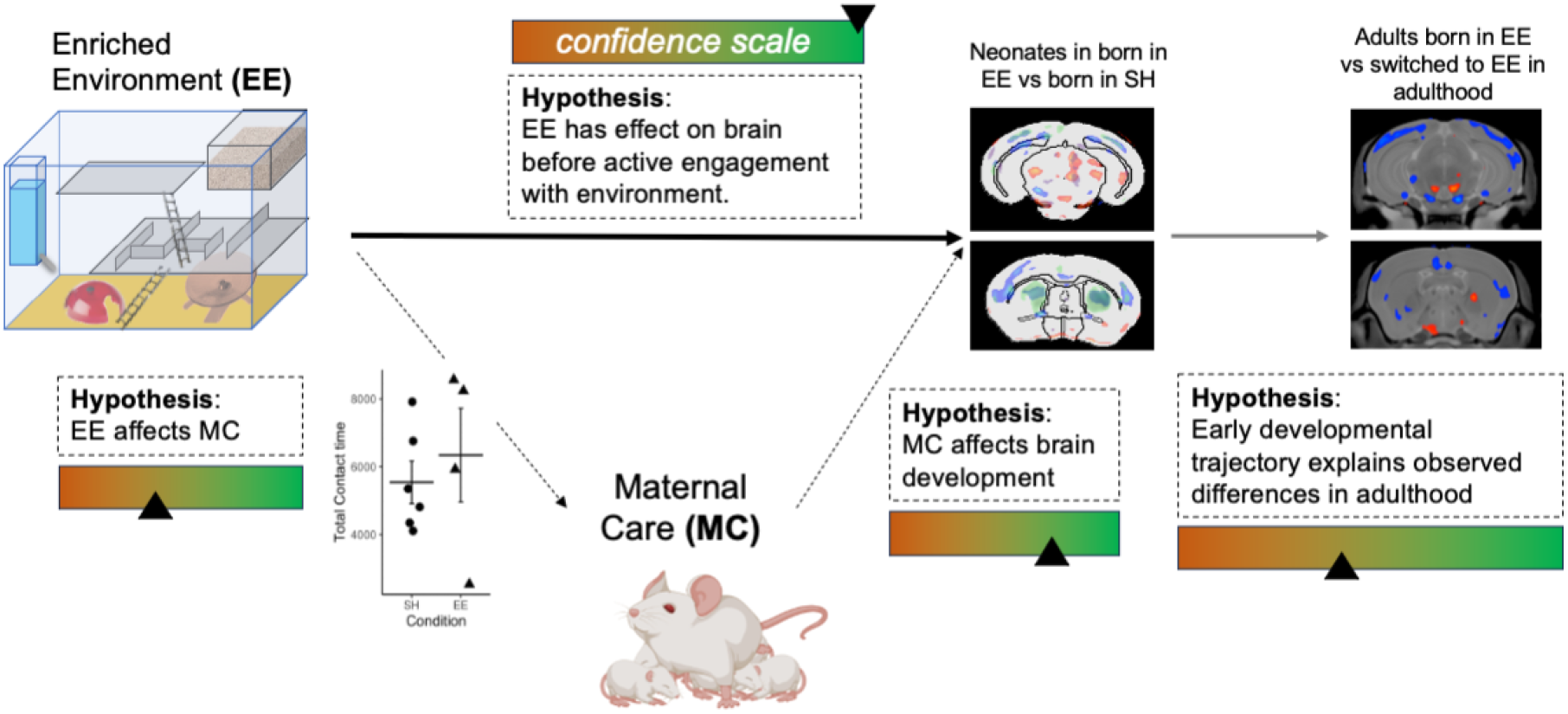
Summary of hypotheses and results.

Our study replicated previous findings showing a large effect of EE on hippocampal volume in adulthood (**Figure 1**, Cohen’s d=1.48 for dataset P and Cohen’s d=1.51 for dataset A)(Scholz et al., 2015), reinforcing that EE is a powerful modulator of plasticity in the adult brain.

As the developing brain exhibits heightened susceptibility to experience and environmental conditions (Reh et al., 2020; Han et al., 2022), we expected a larger effect in adult mice that were raised in an enrichment observable rather than those exposed to it only during adulthood. Additionally, we expected that the timing of environmental stimulation during development would lead to distinct effects on brain structure, as brain development transitions through critical periods of heightened sensitivity in early life. While our observations support the notion of a differential effect of enrichment on brain volume at various developmental stages, the observed sequelae on the adult brain are sparse. The differential effects were observed in cortical areas (auditory, somatomotor, orbital and visual), the piriform cortex and the cerebellar lobules IV-V. These findings suggest that enrichment during early life leads to subtle yet distinct structural changes compared to enrichment introduced only in adulthood, highlighting the impact of developmental timing on experience-dependent plasticity.

To explore the effects of EE in early life we examined the effects of enrichment at postnatal day 7 (P7), before neonates were able to actively explore their surroundings. EE led to significant brain volume changes at P7 (**Figure 2**), with the largest effects observed in the hindbrain and the striatum. Interestingly, EE-induced changes in the hippocampus were focal and consisted of a decrease in apparent volume, as opposed to the increase reported in adulthood. Although the segmentation is coarser at P7 than in adulthood, we could report smaller parietal and occipital lobes apparent volumes, corresponding roughly to the visual areas, the auditory areas and the somatomotor/sensory cortices. These findings highlight the broad effect of EE on whole brain structure during the perinatal period.

Structural MRI is blind to the nature of cellular and molecular mechanisms underlying the changes, but a few insights can be drawn from the literature. At P7, mice’s sensory (vision, audition) and motor systems are immature (eyes and ear canal closed, mobility reduced), and they cannot actively explore their environment. However, early brain development studies focussing on the effect of EE on the visual cortex found that the latter matures faster when neonates are born in an enriched environment (Baroncelli et al., 2010). Indeed, differences in retina maturation are already observed in utero (Sale et al., 2007), and BDNF is more expressed at P7 in the visual cortex (Sale et al., 2004). Moreover, the basic behaviour (motor and auditory reflexes) of neonates is improved from birth for animals born in an EE (Cárdenas et al., 2015). Lastly, a study looking at the hippocampus of neonates from a dam freely exercising during pregnancy showed a stronger cell proliferation in the dentate gyrus of offspring, and most notably, a higher count of astrocytes (Bick-Sander et al., 2006). These studies altogether reinforce our findings regarding early brain structural and microstructural differences in the cortical plate (isocortex and hippocampus) between neonates born in an EE compared to neonates born in SH.

The absence of longitudinal data prevents us from drawing causal conclusions about the volume differences reported in adulthood. The disparity between datasets is also a limiting factor in interpreting the data. Between datasets N and P, dams were transferred to an enriched environment earlier during gestation for N (E13 vs E17), which could matter if in utero effects are strong. Moreover, although effect of litter size was mitigated in datasets N and P, we cannot exclude some variability linked to the partly uncontrolled litter size. These considerations apart, the combination of datasets N, P and A is still indicative. When enriched from birth, animals at P7 (Figure 2A) and adulthood (Suppl. Figure 4, second column) present a relative cortical volume decrease compared to animals raised in SH . Besides those similitudes, most of the volumetric changes observed at P7 are not straightforwardly associated with those observed in adulthood. They may be transient, representing a change in regional brain plasticity and maturation at a stage of extensive brain development (Semple et al., 2013; Tooley et al., 2021; Malave et al., 2022), yet with little to no lasting volume change in this region in later life. It is also possible that the areas highlighted at P7 under EE simply reflect the most plastic regions at this developmental stage. These considerations do not exclude the possibility of lasting functional and microstructural differences of such early life changes, which cannot be measured via conventional volumetric analysis used in this study. Finally, it is possible that these early transient changes in brain structure drive changes in sensory areas in later life, through altered connectivity and neuronal input in early life, or indicate an altered trajectory of brain development with lasting consequences (Tooley et al., 2021).

The neonates’ environment at P7 mostly consists of contact with their mother and siblings. Therefore, a possible explanation for these early changes observed in the brain lies in the nature of maternal behaviour, which has a critical influence on development (Champagne et al., 2003; Meaney et al., 2007). Likewise, our study suggests that maternal behaviour affects early postnatal brain development (**Figure 3**). Furthermore, the significant overlap between brain volume changes observed due to EE and maternal behaviour (R=0.79), together with the prominent role of maternal contact at the age of P7, suggest that the effect of EE may partially be mediated by changes in maternal behaviour (**Figure 5**). However, evidence for a link between EE and maternal behaviour, supported by a range of studies (Sparling et al., 2020), is limited by measurement methods and insufficient power in the current study. Nonetheless, our observations reinforce the idea that maternal behaviour is a potential route by which EE may influence brain development before the neonates can actively explore the environment.

Overall, this study provides novel insights into the effect of perinatal environmental enrichment during the early postnatal period and later developmental stages. Future studies should longitudinally explore early life changes to better understand how they affect developmental trajectory. Further, careful choice of litter size and dam number can increase statistical power to assess the effect of maternal behaviour. Cross-fostering experiments can help disentangle *in utero* from postnatal effects. Finally, investigating the micro- and macrostructural factors underlying the observed volumetric changes should be probed to gain further insight into the mechanisms by which perinatal enrichment affects early brain development. Typically, microscopy or advanced MRI methods can unravel the microscopic nature (e.g. spine density, dendritic branching, gliosis, angiogenesis etc) of the volumetric changes observed. Other in vivo (e.g. dMRI, fMRI) or ex vivo (e.g. spatial transcriptomics) imaging modalities can help explore further the connectivity and relationships between EE highlighted brain regions.

## Methods

### Animals

Details about animals of each dataset can be found in the section **Datasets** below. CD-1 stock mice and breeding pairs were acquired from the active colony at the Toronto Centre for Phenogenomics (*Toronto, Ontario, Canada*) and were maintained on a 12-hour light/dark cycle with *ab libitum* access to food (low fat diet, *Harlan Teklad #2914*) and water. Mice were housed either in standard housing cages (SH) or within an enriched environment (EE) as described in section **Enrichment setup** below. All animal experiments were approved by the animal ethics committee of the Toronto Centre for Phenogenomics.

### Enrichment setup

The cage acted on all component susceptible to increase hippocampal volume (maze: navigation skills, free running wheel: exercise, housing dome: environmental enrichment). The control cage had the same size but no 3D maze, no wheel, no housing dome. Social enrichment was kept constant (7 mice per condition). The setup was adapted from (Scholz et al., 2015) and is briefly described below.

The cage of the enriched environment was a modified version of the Double Decker Rat IVC GreenLine cages (Tecniplast, Italy) with dimensions 462mm wide × 403mm deep × 404mm in height and a floor area of 1862 cm^2^. The cage was retrofitted from its standard configuration by removing the second-level floor board and replacing it with a multi-level maze consisting of interlocking polycarbonate walls, ceilings/floors and pipes . The mazes pieces were designed in such a way that they can be easily rearranged to change the spatial layout of the maze and create new pathways for the mice to navigate. At every cage cleaning we altered the maze according to a sequence of pre-specified maze layouts . Water bottles were fitted with long spouts to allow mice to drink from the ground level. The food hopper was fitted with a stainless-steel lid and placed on the top level of the maze to maximize the distance between food and water. Grain and bedding material were added to the floors on each maze level along with a dome and running wheel to provide additional enrichment. This 3-dimensional maze allows mice to exhibit their natural behaviour of navigating through a burrow.

### Datasets

Three different datasets were used in this study, with different timings of exposure to enriched environment.

**N:** “P7 perinatal enrichment dataset”: Five CD1 dams were mated in standard housing. At E13, two dams were transferred to an enriched environment and three were kept in standard housing. The EE cage was slightly modified to allow for visualization and assessments of maternal behaviour. As an added benefit, this modification leads to an open nesting area that has similar dimensions to the SH cage. The number and sex of neonates in the litter were determined on P0 (Table 3. Litter size and sex distribution Suppl. Table 1). On P3 and P4, maternal behaviour was assessed (see Table 4 for details). Neonates from these first litters were perfused at P7 for ex vivo MRI (see **MRI Acquisitions**). Dams were placed back in standard housing. Six weeks later, the same dams were mated again, and at E13, the dams were transferred to the same environments than before, except for 2 dams that were switched environment (EE > SH and SH > EE). Only 11 neonates per litter were kept at birth for these second litters to mitigate litter size effects. Neonates were perfused at P7 for ex vivo MRI (see **MRI Acquisitions**). 119 neonates were born (Supplementary Table 1); 1 was not scanned, and a further 4 were excluded due to poor image quality, yielding a final sample of 114 neonates (49 EE: 26 females, 23 males; 65 SH: 28 females, 37 males) used for analysis. The behaviour of dams was assessed when neonates were P3 and P4 (see **Behaviour**). The summary of the experimental design is shown in **Suppl. Figure 5A**. This design provides control for natural variations in maternal behaviour and genetic differences that may influence neuroanatomy by using the same breeding pair for multiple litters and exposing them to one housing condition followed by the other.

**P:** “perinatal enrichment dataset”: at E17, two pregnant CD1 dams were transferred to an enriched environment and two were kept in standard housing. At weaning neonates were kept in the same environment but males and females were separated and neonates from the 2 litters were mixed to alleviate litter effects. All mice were perfused at P43 for ex vivo MRI (see **MRI Acquisitions**). The dataset contains 49 brains, with 24 animals born in an EE (13 females), and 25 born in SH (14 females).

**A:** “adult enrichment dataset”: 27 CD1 mice (25 + 2 outliers) male mice were included. 13 (2 outliers) were transferred in an enriched environment at P53 for 43 days. 14 stayed in their standard environment. All mice were perfused at P96 for ex vivo MRI (see **MRI Acquisitions**).

### Perfusions

Dataset N: In preparation for ex vivo imaging, P7 neonates were separated from their mothers, immediately anesthetised (150 mg/kg of ketamine and 10 mg/kg of xylazine) and perfused through the heart with 10 mL of heparin diluted in phosphate-buffered saline (PBS), followed by 10 mL of 4% paraformaldehyde (PFA) diluted in PBS. Following decapitation, skulls containing the brain were post-fixed for 24 hours at 4°C in a 4% solution of PFA. The skulls were then placed into a solution of 0.01% sodium azide in PBS at 4°C until the time of scanning.

Dataset P and A: In preparation for ex vivo imaging, mice at P43 (dataset P) or P96 (dataset A) were deeply anesthetised with a ketamine-xylazine mixture. They were then perfused through the left ventricle with 30 mL of phosphate-buffered saline (PBS) and 1 μL/mL heparin (1000 USP units/mL). This was followed by infusion with 30 mL of 4% paraformaldehyde (PFA) in PBS for fixation as described in (Cahill et al., 2012).

### Behaviour

#### Maternal Behaviour assessments on P3-4

Before the assessment of maternal behaviour, cages were moved back and forth between the housing facility and the testing room to habituate the dams to the movement. Assessment of maternal behaviour took place on P3 for 35 mins (between 10 am and 12 pm) and on P4 for four 30 mins sessions spread out between 9 am and 5 pm (total of 2 hours and 35 mins per cage). Each cage was observed for the 30 min session and then the next cage was assessed in the sequence. This protocol for assessing maternal behaviour was based on Myers et al. (1989) and Champagne et al., (2003) with an emphasis on active nursing postures and grooming bouts. Suppl. Table 3 shows which behaviours were scored and how they were collapsed for analysis. During the sessions, the observer noted the onset and offset of each behaviour and recorded the time providing a direct measure of duration (seconds). Each session was video recorded.

#### MRI scanning

Datasets A and P: 3D fast spin-echo sequence, TR = 2000 ms, echo train length=6, TE_eff_=42 ms, field-of-view (FOV)=25 × 28 × 14 mm and matrix size=450 × 504 × 250 (i.e., a 56µm isotropic resolution), for a total imaging time of approximately 12 hours. No contrast agent was used.

Dataset N: Brains of one-week-old neonates were imaged using a 3D diffusion-weighted fast spin echo sequence: TR= 350 ms, Echo Train Length = 6, first TE= 30 ms, subsequent TEs = 6 ms, 1 average, FOV = 2.5 x 1.4 x 1.4 cm^3^. A cylindrical acquisition was used to acquire 77% of the k-space data from a matrix of 324 x 180 x 180, which yielded an image with 78 μm isotropic voxels. Thirty images corresponding to 30 directions at b=1917 s/mm^2^ were acquired and averaged. The total imaging time was 14 hours. No contrast agent was used, as the diffusion-weighting provided the contrast necessary for analysis.

### Analysis

#### Image registration

Image registration allows the quantification of anatomical differences between images. For a group of images, this procedure results in a transformation that maps every point in one image to corresponding points in the other images. Thus, the differences between the images are captured by this transformation. Our procedure for image registration is composed of an affine registration, followed by a series of non-affine registrations. The affine registration applies global translation, rotation, scaling, and shearing to align images. Information regarding global deformations (i.e. the overall brain sizes) are stored in these transformation models. The non-affine registration creates a vector field that maps every point in one image to another and provides information about localized deformation.

Data were analysed using the MBM Pydpider toolkit (Friedel et al., 2014) with a MAGeT registration for segmentation. The MRI atlas (Richards et al., 2011; Ullmann et al., 2013; Steadman et al., 2014; Beera et al., 2018; Qiu et al., 2018) was used for dataset A and P. For the P7 data, a resampled and simplified version MRI atlas of the same atlas was applied. Group-wise registration was performed on datasets A and P together and on dataset N independently. Regional volumes were extracted from the individual LSQ6 resampled atlases with the RMINC tools. Hierarchical anatomical trees to visualise overall regional changes.

#### Statistical analysis

For voxel-wise analysis, linear models were applied voxel-wise and multiple comparisons were corrected with an FDR threshold. The mincLm function from the RMINC package in R was used. The regressors were the following:

- Dataset A and P, to look at the effect of housing: housing condition (SH or EE), group (A or P), sex (F or M)

- Dataset A and P, to look at the differential effect of housing: housing condition*group, sex

- Dataset N, to look at the effect of housing: housing condition (SH or EE), sex (F or M), litter order (first litter or second litter), litter size (number of neonates in the litter).

Similar linear models were also applied to the regional volumes (via the hierarchical volumetric organisation with the hanatLm function in RMINC). This enabled us to identify the regions of interest (**Figure 1D** and **Figure 2C**):

- Datasets A and P: Regional volumes were normalised to the total brain volume. For the z-scored regional volumes, normalised data are zeroed to the mean of the standard housing condition in their age group, because mice continue to develop slightly between P43 (perfusion age of dataset P) and P96 (perfusion age of dataset A), and relative regional volumes differ between these 2 ages. The z-score is calculated using the mean and the standard deviation is shared over datasets P and A. Note that only male volumes are shown in **Figure 1D** to address the sex discrepancy during development (for dataset P, since dataset A does not contain females).

- Dataset N: For the z-scored regional volumes, normalised data are zeroed to the mean of the standard housing condition in their age group. The z-score is calculated using the mean and the standard deviation is shared over datasets P and A. Note that only male volumes are shown in **Figure 2C** to address the sex discrepancy during development (for dataset P).

#### ROI data extraction

To extract volumetric changes induced by EE from regions of interest (ROI) in the N dataset and compare them to the effect of maternal contact time (**Figure 3F**, Suppl. **Figure 5F**), t-stats maps from the effect of EE on brain structure were thresholded at the 1% FDR level (t=3.2) and binarized. ROI masks were then created from these binarized t-stats maps for both Striatum and Brainstem (relying on the atlas anatomical landmarks). These regions were picked because they were significantly affected by EE, in a large and bilateral manner. Mean relative Jacobian values were extracted from these ROI. The litter size effect was controlled using linear regression. From the resulting residual values, z-score were calculated and compared against maternal contact time.

#### Mediation analyses of housing effects on brain structure

To test whether maternal care mediated the effects of postnatal housing condition (standard vs. enriched) on brain structure, we performed mediation analyses at both the region-of-interest (ROI) and voxel-wise levels. Housing condition was modelled as a categorical independent variable, maternal contact behaviour as a continuous mediator, and litter size was included as a covariate in all analyses.

### ROI-based mediation analysis

Initial analyses were conducted on predefined anatomical regions of interest (ROI) to establish the statistical presence, direction, and magnitude of mediation effects in summary brain measures. Mediation analyses followed a causal mediation framework implemented in R, with linear regression models specified for the mediator (maternal contact behaviour) and the outcome (regional brain volume). Average causal mediation effects (ACME), average direct effects (ADE), total effects, and proportions mediated were estimated using nonparametric bootstrap inference with 3,000 resamples.

This frequentist approach assumes independence of observations and therefore does not explicitly model the nested structure of pups within mothers. It was used to provide an interpretable baseline assessment of mediation effects at the regional level and to enable direct comparison with voxel-wise analyses.

To account for the hierarchical structure of the data arising from pups nested within individual mothers, complementary Bayesian multilevel mediation models were fitted. These models included random intercepts for mother identity and were implemented using the brms package in R. Posterior distributions of indirect, direct, and total effects were derived by combining posterior samples of the relevant path coefficients, and uncertainty was summarised using medians and 95% credible intervals.

Although these Bayesian models were not treated as primary outcome analyses, they yielded results that were qualitatively consistent with the frequentist ROI findings. Specifically, housing showed a strong overall association with striatal volume (posterior median = −1.08, 95% credible interval: −1.77 to −0.18), with evidence that a substantial proportion of this effect was mediated by maternal contact behaviour (median proportion mediated = 47%). The posterior distribution of the indirect (mediated) effect was centred below zero (median = −0.52), with a credible interval that narrowly excluded zero (−1.26 to −0.02), indicating moderate support for a mediation pathway. The direct effect of housing after accounting for maternal contact was more uncertain (median = −0.54, 95% credible interval: −1.27 to 0.28), suggesting that while maternal care accounted for a meaningful portion of the housing effect, additional non-mediated pathways may also contribute. Given the limited number of mothers, these Bayesian estimates were interpreted cautiously and used primarily to assess robustness to model assumptions.

### Voxel-wise mediation analysis

To spatially characterise mediation effects across the brain, the mediation framework was extended to a voxel-wise analysis. Using the same model structure and covariate adjustment as in the ROI analyses, mediation models were fitted independently at each voxel within a predefined brain mask. For computational feasibility, voxel-wise analyses were performed using a frequentist mediation approach.

Voxel-wise mediation effects - including ACME, ADE, total effect, and proportion mediated - were estimated using quasi-Bayesian simulation based on the asymptotic normality of the regression coefficients, without nonparametric bootstrap resampling, with 3,000 simulation draws per voxel. Whole-brain analyses were implemented using the RMINC framework, enabling efficient parallel computation. Voxel-wise mediation maps were used to visualise the spatial distribution, directionality, and relative strength of direct and indirect housing effects on brain structure and were interpreted as exploratory and descriptive.

## Data and code availability statement

The data and code used in this study will be made publicly available upon publication. Upon acceptance, they will be deposited in GitHub and Zenodo and assigned a DOI for permanent access. Prior to publication, data and code may be shared upon reasonable request.

## SUPPLEMENTARY

**Suppl. Figure 1:**
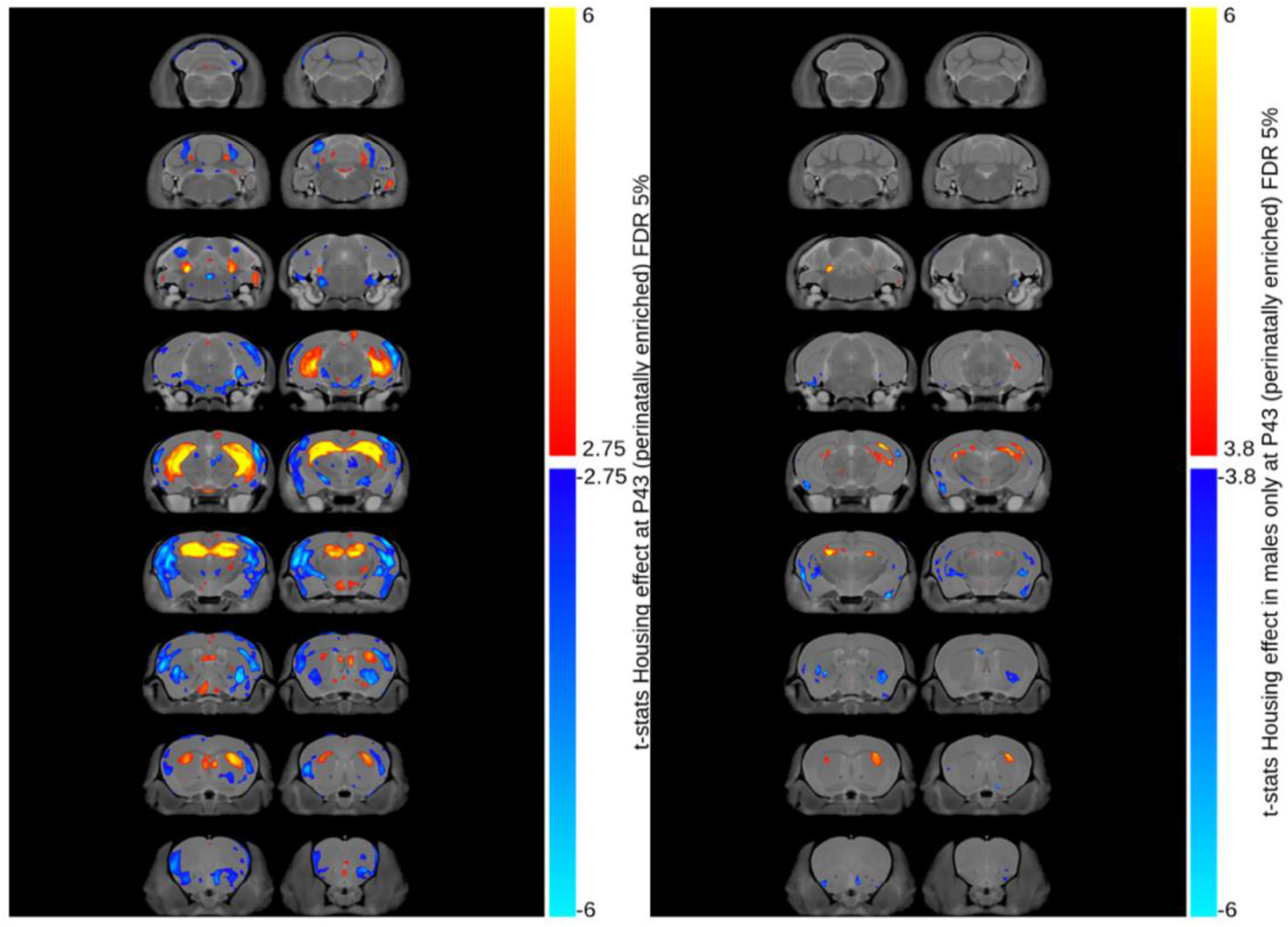
Males only in P-data. A voxel-wise linear model was applied to the Jacobians computed after linear co-registration (hence corrected for individual brain volume variation) coming from datasets P (left column) and A (right column). The regressor is housing condition.

**Suppl. Figure 2:**
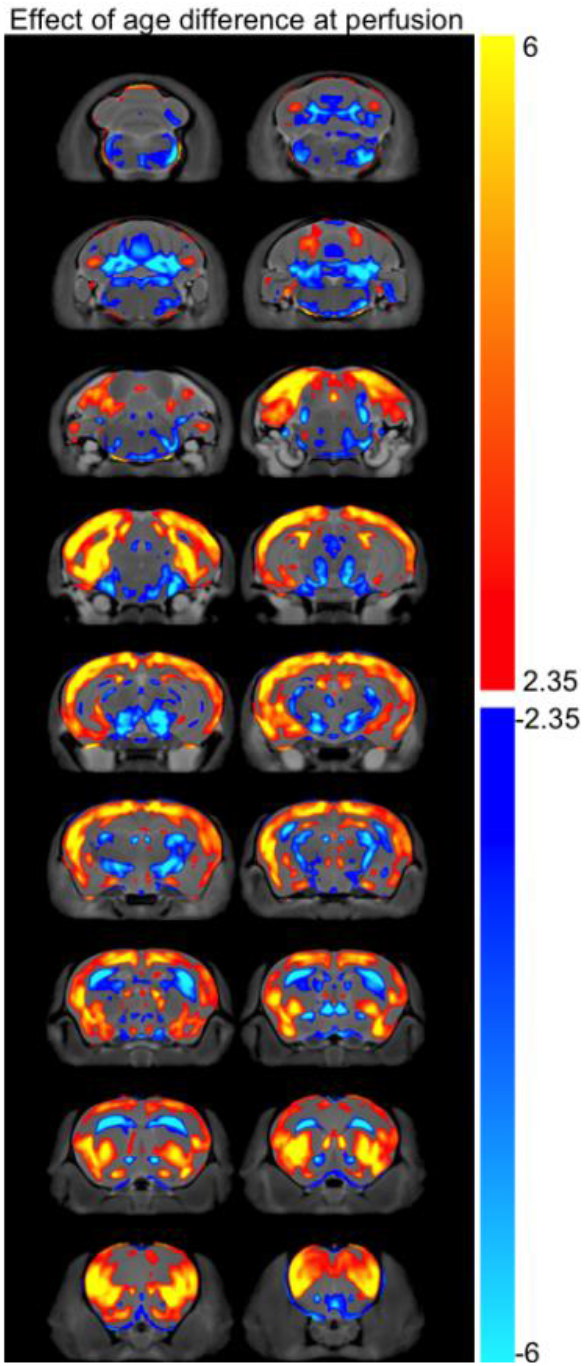
In addition to Figure 1C. Effect of age at perfusion (P43 vs P96). A voxel-wise linear model was applied to the Jacobians computed after linear co-registration (hence corrected for individual brain volume variation) coming from datasets P and A. The regressor are housing condition, age and sex, and the effect of age is shown.

**Suppl. Figure 3:**
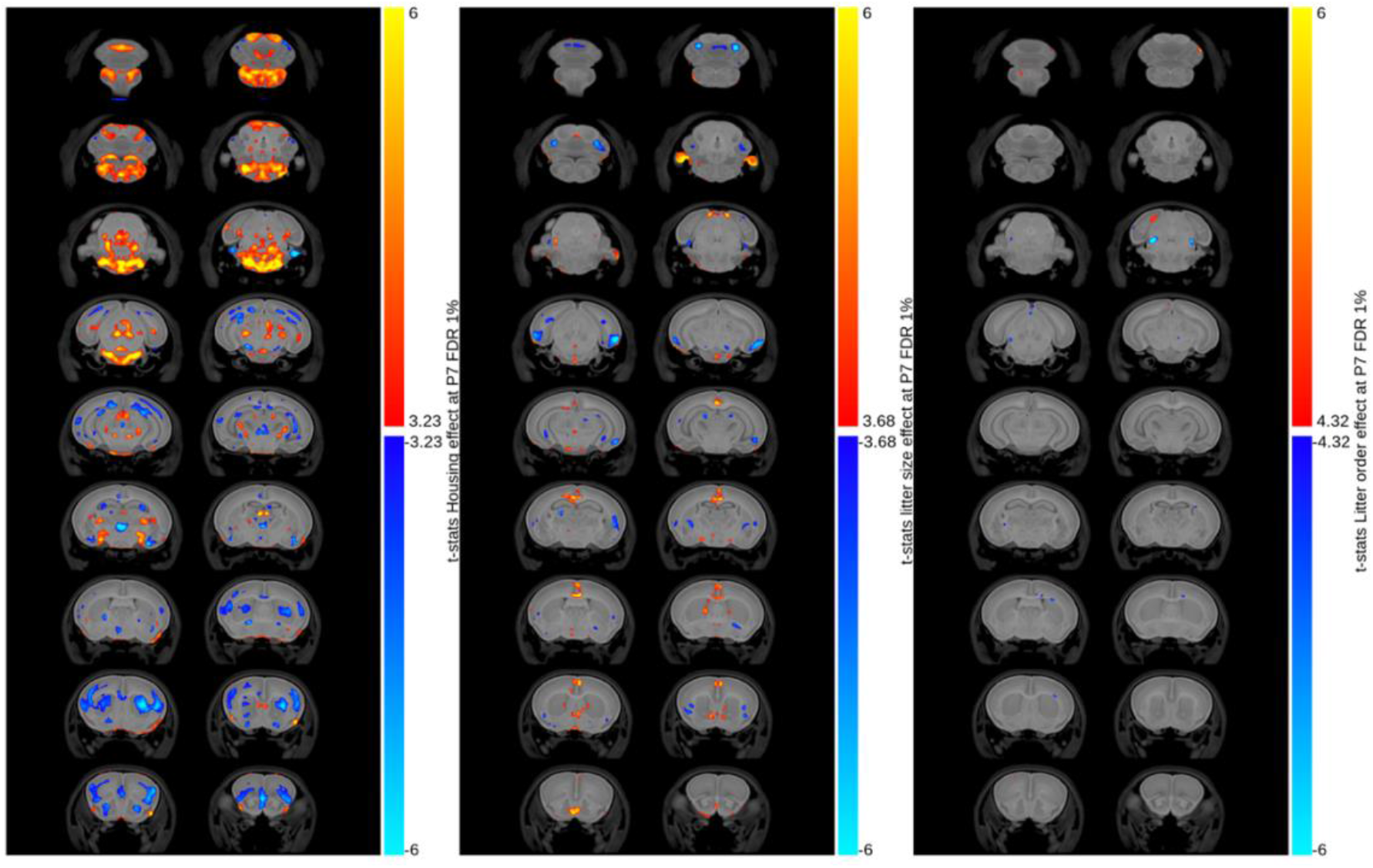
Effect of Litter Size and Litter order at P7 (N). A linear model was applied to the Jacobians computed after linear co-registration (hence corrected for individual overall brain volume variation), with the following housing condition, litter order, litter size and sex as regressors. The effect of housing condition is shown with a FDR threshold at 1% (left column). The effect of litter size is shown with a FDR threshold at 1% (middle column). The effect of litter order is shown with a FDR threshold at 1% (right column).

**Suppl. Figure 4:**
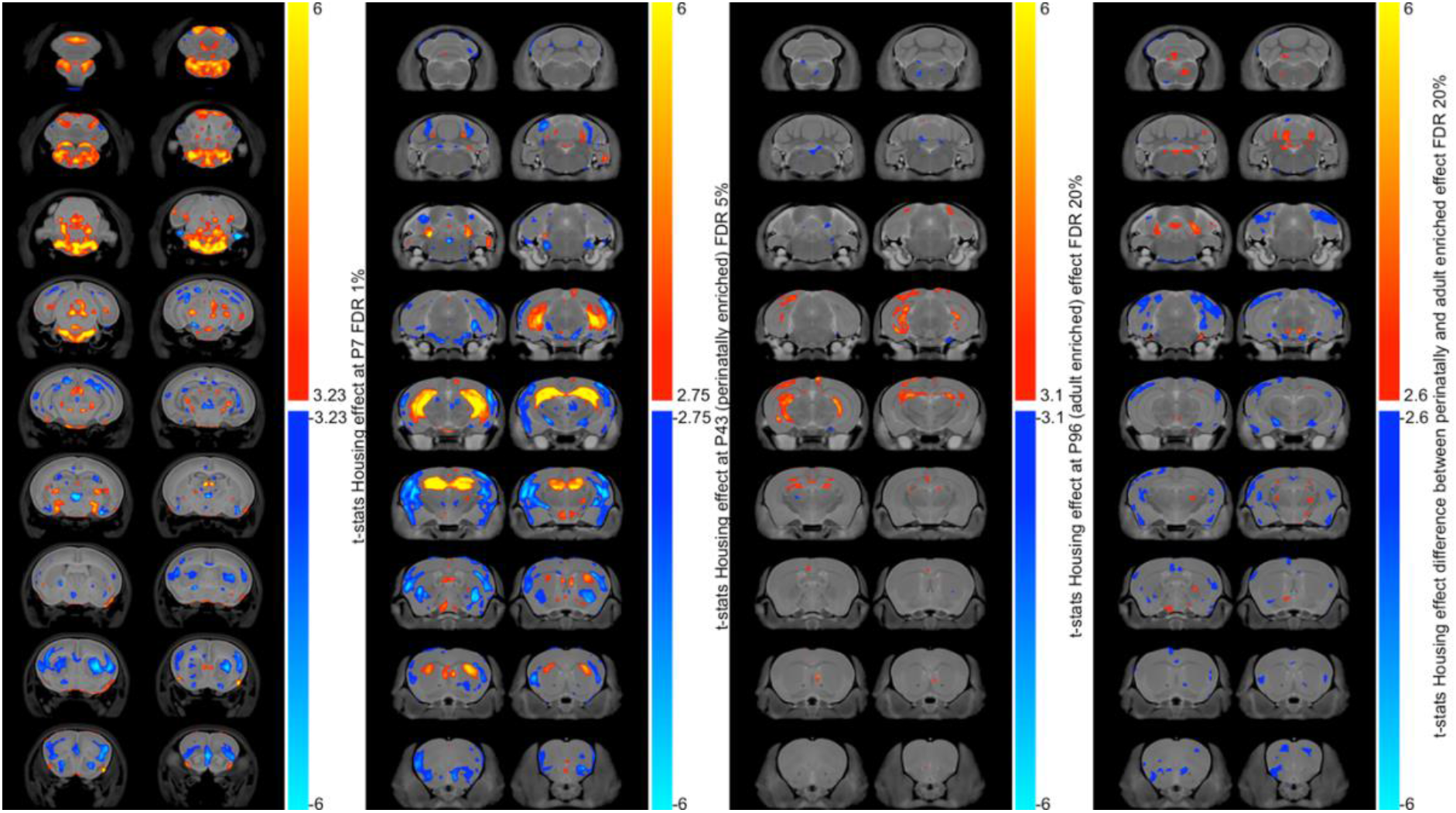
(N-P-A-P|A) housing comparison. Visual comparison of the impact of housing condition on brain volume for different datasets. L<u>inear models were applied to the Jacobians computed after linear co-registration (hence corrected for individual overall brain volume variation). First column, regressors:</u> housing condition, litter order, litter size and sex; dataset: N. Second column, regressors: housing condition and sex; dataset: P. Third column, regressors: housing condition and sex; dataset: A. Fourth column: housing, age and sex, and the interaction term housing*age; dataset: A+P.

**Suppl. Figure 5:**
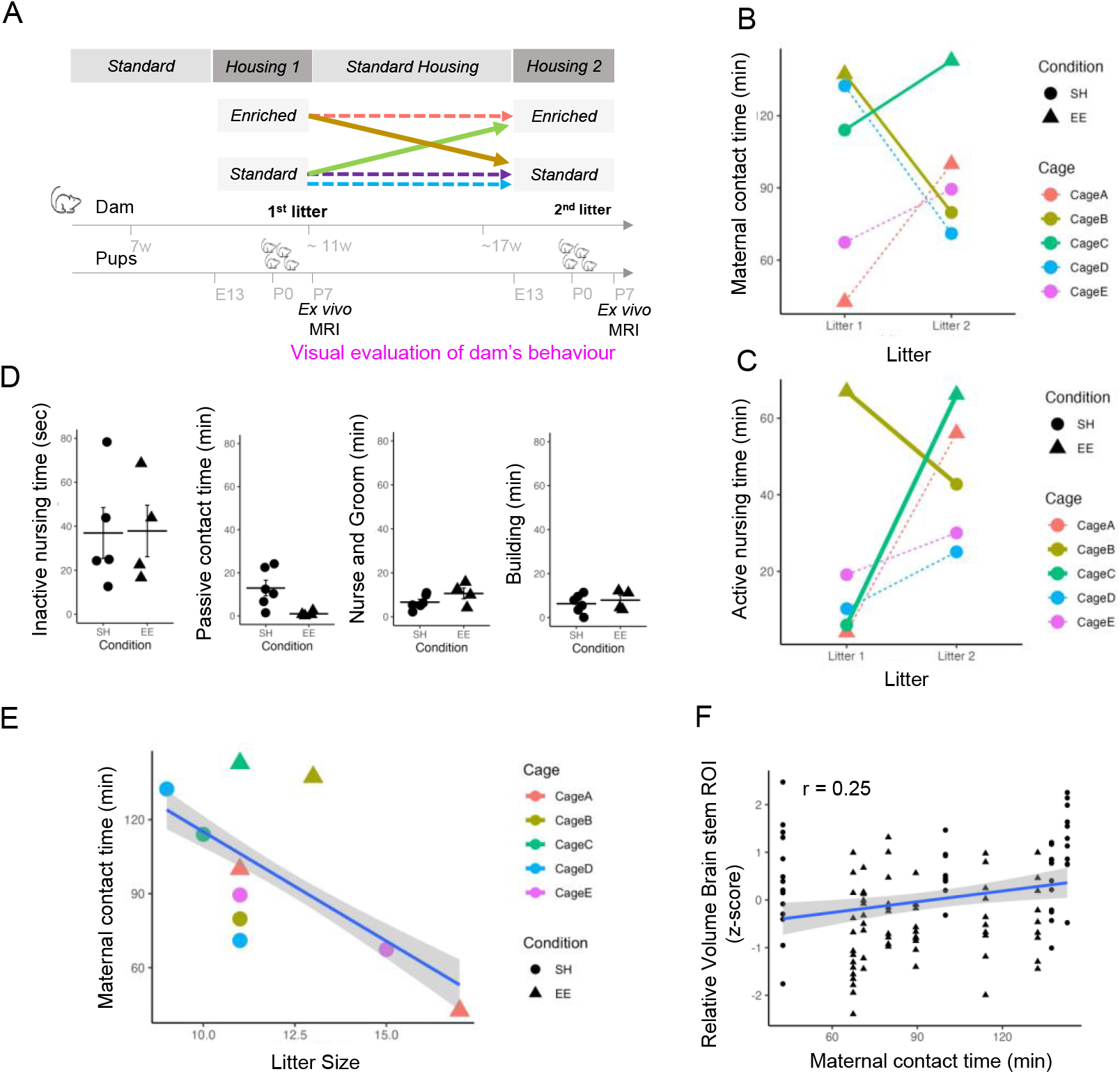
Maternal care, perinatal enrichment and volumetric changes in P7 brains. **(A)** Experimental design for assessing maternal care and brain stricture at P7. **(B)** Quantity of observed maternal contact time per housing (Triangle is EE) for the 1^st^ and 2^nd^ Litter. **(C)** Quantity of observed active nursing time per housing for the 1^st^ and 2^nd^ Litter for each dam. **D.** Quantification of additional maternal contact behaviors per condition **E.** Relationship between Total contact time of mums with the neonates and the number of neonates in the litter. **F.** Volumetric change in Brain stem ROI plotted against total contact time of the damn during the observed time period (r = 0.25)

**Suppl. Table 1:** Overview of subjects numbers in different dams (Cage A-2) and Litter (1 or 2) in the sample of P7 old pubs scanned ex vivo.

| Breeding Pair | Litter | Housing Condition | Female pups (n=) | Male pups (n=) | Total pups (n=) |
| --- | --- | --- | --- | --- | --- |
| Cage A | 1 | Enriched (EE) | 9 | 8 | 17 |
| Cage A | 2 | Enriched (EE) | 6 | 5 | 11 |
| Cage B | 1 | Enriched (EE) | 9 | 4 | 13 |
| Cage B | 2 | Standard (SH) | 6 | 5 | 11 |
| Cage C | 1 | Standard (SH) | 5 | 5 | 10 |
| Cage C | 2 | Enriched (EE) | 5 | 6 | 11 |
| Cage D | 1 | Standard (SH) | 2 | 7 | 9 |
| Cage D | 2 | Standard (SH) | 3 | 8 | 11 |
| Cage E | 1 | Standard (SH) | 7 | 8 | 15 |
| Cage E | 2 | Standard (SH) | 6 | 5 | 11 |
| TOTAL |  |  | 58 | 61 | 119 |

**Suppl. Table 2:**
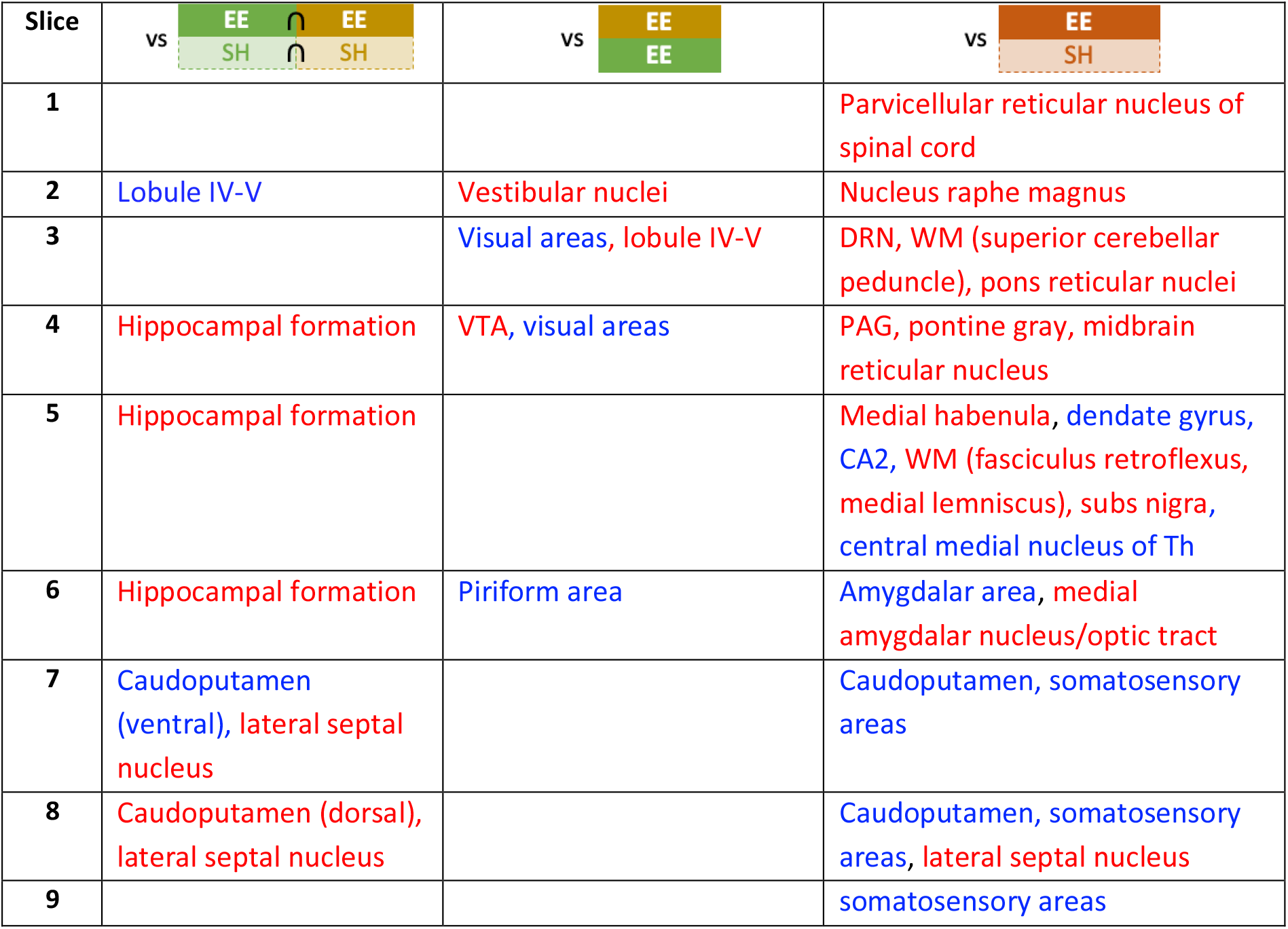
Volumetric changes due to enrichment. Slice number corresponds to presentation in Figure 1 and Figure 2. Text colour indicates direction of change (Increase:red; decrease: blue)

**Suppl. Table 3:** Maternal behaviour categories, code and description of behaviour used for categorization.

| Category | Code | Description of Behaviour |
| --- | --- | --- |
| **Building** | B | Nesting Building. Retrieving bedding and grain from around the cage and placing it in the vicinity of the nest. No contact with the pups. |
|  | CB | Contact and building. Contact with the pups while building and working on the nest. |
| **Eating/Drinking/**<br>**Sleeping** | X | Sleeping away from pups and nest. |
|  | SX | Self-grooming alone. |
|  | EX | Eating alone. |
|  | DX | Drinking alone. |
| **Nest Contact** | C | Contact. Physical contact with the pups that is not grooming, nursing or building. |
|  | CE | Contact eating. When in contact with the pups she may be eating: food pellets and/or grain. |
|  | CS | Contact self-grooming. When she is in contact with the pups and is grooming herself. Distinct from self-grooming that occurs in conjunction with pup grooming. |
| **Passive Nursing** | P | Passive nursing pose. Dam is lying on her side to allow the pups to nurse. She is putting minimal effort into the act and is usually sleeping. |
|  | N1 | Low nursing pose. When the dam is lying on top of the pups putting minimal effort into the nursing pose. Not using her legs to support her body weight. 'Superman pose'. |
| **Active Nursing** | NE | Nursing and eating. |
|  | N2 | Medium nursing pose. When the dam is on top of the pups forming a slight arch in her back. She is putting weight on her legs to support the arch and permit the movement of pups under her belly. |
|  | N3 | High nursing pose. When the dam is on top of the pups putting maximal effort into the nursing pose and forming an arch in her back. Ventrums can be seen permitting the movement of pups under her belly. |
| **Nursing/Grooming** | G | Grooming. Grooming pups without nursing. |
|  | N1G | Grooming with low nursing pose. |
|  | N2G | Grooming with medium nursing pose. |
|  | N3G | Grooming with high nursing pose. |
| **Contact** | - | Any behaviour in which the dam was in physical contact with pups. Includes the following behaviours (codes): C, CB, CE, CS, P, NE, N1, N2, N3, G, N1G, N2G, N3G. |
| **Nursing** | - | Any nursing behaviour regardless of posture. Includes the following behaviours (codes): P, NE, N1, N2, N3, N1G, N2G, N3G. |

